# Lessons from the meiotic recombination landscape of the ZMM deficient budding yeast *Lachancea waltii*

**DOI:** 10.1101/2021.12.13.472358

**Authors:** Fabien Dutreux, Abhishek Dutta, Emilien Peltier, Sabrina Bibi-Triki, Anne Friedrich, Bertrand Llorente, Joseph Schacherer

## Abstract

Meiotic recombination has been deeply characterized in a few model species only, notably in the budding yeast *Saccharomyces cerevisiae*. Interestingly, most members of the ZMM pathway that implements meiotic crossover interference in *S. cerevisiae* have been lost in *Lachancea* yeast species after the divergence of *Lachancea kluyveri* from the rest of the clade. This suggests major differences in the control of crossover distribution. After investigating meiosis in *L. kluyveri*, we determined the meiotic recombination landscape of *Lachancea waltii* and identified several characteristics that should help understand better the underlying mechanisms. Such characteristics include systematic regions of loss of heterozygosity (LOH) in *L. waltii* hybrids, compatible with dysregulated Spo11-mediated DNA double strand breaks (DSB) independently of meiosis. They include a higher recombination rate in *L. waltii* than in *L. kluyveri* despite the lack of multiple ZMM pro-crossover factors. *L. waltii* exhibits an elevated frequency of zero-crossover bivalents as *L. kluyveri* but opposite to *S. cerevisiae. L. waltii* gene conversion tracts lengths are comparable to those observed in *S. cerevisiae* and shorter than in *L. kluyveri* despite the lack of Mlh2, a factor limiting conversion tracts size in *S. cerevisiae. L. waltii* recombination hotspots are not shared with either *S. cerevisiae* or *L. kluyveri*, showing that meiotic recombination hotspots can evolve at a rather limited evolutionary scale within budding yeasts. Finally, in line with the loss of several ZMM genes, we found only residual crossover interference in *L. waltii* likely coming from the modest interference existing between recombination precursors.

**Significance statement:** Studying non-model species is relevant to understand better biological processes by shedding light on their evolutionary variations. Here we chose the non-model budding yeast *Lachancea waltii* to study meiotic recombination. In sexually reproducing organisms, meiotic recombination shuffles parental genetic combinations notably by crossovers that cluster in hotspots at the population level. We found remarkable variations compared to both the canonical *Saccharomyces cerevisiae* model and another close relative *Lachancea kluyveri*. Such variations notably include the loss in *L. waltii* of a layer of regulation of crossover distribution that is otherwise conserved in budding yeasts and mammals. They also include the lack of conservation of crossover hotspots across the *Lachancea* species while crossover hotspots are remarkably stable across the *Saccharomyces* species.

**Highlights:**

- Extensive LOH events in *L. waltii* intraspecific hybrids
- No conservation of the recombination hotspots across the *Lachancea* genus
- Reduced but not suppressed crossover interference in the absence of the ZMM pathway
- Similar gene conversion tract lengths in *L. waltii, S. cerevisiae*, and *L. kluyveri* despite the lack of *MLH2* in *L. waltii*

## Introduction

Homologous recombination is a ubiquitous DNA repair process that generates crossover and noncrossover recombinants (Mehta and Haber. 2014). In most species, crossovers resulting from meiotic recombination are essential for accurate homologous chromosome segregation at the first meiotic division and hence fertility (Petronczki et al. 2003). In addition, by promoting shuffling of parental alleles, both crossover and noncrossover recombinants promote diversity (Stapley et al. 2017a). Meiotic recombination is, therefore, under strong selective constraint.

Meiotic recombination results from repairing programmed DNA double strand breaks (DSBs) made at leptotene by a multi-protein complex. The catalytic subunit is the topoisomerase II-like trans-esterase Spo11 protein (Lam and Keeney, 2014; Robert et al. 2016). DSB formation is highly regulated. In many organisms, it eventually results in DSB hotspots in nucleosome-depleted regions within loops of chromatin generated by the compaction of the meiotic chromosomes. DSB hotspots and subsequent recombination hotspots are frequently enriched in gene promoters and other functional elements in organisms lacking the PRDM9 methyl transferase (Auton et al. 2013; Brick et al. 2012; Choi and Henderson 2015; Lam and Keeney 2015; Pan et al. 2011; Singhal et al. 2015). When PRDM9 is present, DSB and recombination hotspots are determined by its sequence specific DNA binding provided by a rapidly evolving minisatellite encoding an array of zinc fingers (Baudat et al. 2010; Diagouraga et al. 2018; Myers et al. 2010; Parvanov et al. 2017). Hence, PRDM9 determined DSB and recombination hotspots are particularly fast evolving, unlike those determined by functional elements that are more stable across evolution (Grey et al. 2018; Lam and Keeney, 2015; Singhal et al. 2015; Tsai et al. 2010).

In addition to the determining effect of DSBs, recombination events distribution along chromosomes undergoes other selective constraints, notably because some configurations are at risk. Due to its repetitive nature, DSB formation and recombination are at risk in the rDNA locus (Nambiar et al. 2019). Crossover formation near centromeres may threaten chromosome segregation during meiotic divisions I and II and are therefore repressed, primarily by preventing DSB formation nearby (Nambiar and Smith 2016). In different yeasts and fungi, recombination is prevented near the mating-type locus, sometimes over several hundreds of kilobases corresponding to an entire chromosome arm (Hartmann et al. 2021). This may be linked to the risk of rendering the mating type locus homozygous and hence perturbing allele frequency of the two mating types within the population. Last but not least, while a single crossover combined with sister chromatid cohesion allows proper tension to be generated between homologous chromosomes at meiosis I, the presence of a second crossover at the vicinity of the first one would likely prevent such tension from being generated due to restrained cohesion between sister chromatids, hence promoting chromosome missegregation at meiosis I. Crossovers, therefore, tend not to form at the vicinity of one another as a result of a mechanism called crossover interference (Berchowitz and Copenhaver 2010). The molecular mechanism of crossover interference is so far unknown. In *S. cerevisiae*, it relies on topoisomerase II (Zhang et al. 2014) and seems independent of the synaptonemal complex. However, SC central region proteins are essential for imposing crossover interference in *Caenorhabditis elegans* and in *Arabidopsis thaliana*, suggesting the existence of different layers of crossover control which contributions vary between species (Libuda et al. 2013; Capilla-Pérez et al. 2021; France et al. 2021). The strength of crossover interference also varies among organisms. Only one crossover per chromosome is formed in *C. elegans* due to a crossover interference strength that exceeds the size of the chromosomes (Libuda et al. 2013). On the contrary, crossover interference hardly spreads above 100 kb in *Saccharomyces cerevisiae* (Anderson et al. 2015). In *S. cerevisiae*, interfering crossovers, also named type I crossovers, are generated through the so-called ZMM pathway, which historically comprises Zip1, Zip2, Zip3, Zip4, Spo16, Msh4, Msh5 and Mer3 (Börner et al. 2004). The Pph3 phosphatase subunit and the proteasome subunit Pre9 are more recently identified members of the ZMM pathway (Pyatnitskaya et al. 2019). The ZMM pathway relies on the nuclease activity of the Mlh1-Mlh3 heterodimer for resolution of recombination intermediates exclusively into crossovers (Allers and Lichten 2001; Cannavo et al., 2020; Hunter and Kleckner 2001; Kulkarni et al. 2020; Nishant et al. 2008). There are also non-interfering crossovers or type II crossovers in *S. cerevisiae* (Santos et al. 2003). They are less abundant than type I crossovers and are formed independently of the ZMM proteins through the resolution of recombination intermediates by the structure specific nucleases Mus81, Yen1 and Slx1 (De Muyt et al. 2012; Zakharyevich et al. 2012). Remarkably, while the ZMM proteins are conserved from *S. cerevisiae* to mammals, they have been lost several times independently during evolution. They are absent from *S. pombe* and from different species of the Saccharomycotina subphylum. They have been lost in *Eremothecium gossypii* after its divergence from *Eremothecium cymbalariae* (Wendland and Walther 2011). They have been partially or completely lost in the *Candida* clade (Clark et al. 2013). And finally, all the ZMM genes except *ZIP1* and *MER3* have been lost in the *Lachancea* clade with the exception of *Lachancea kluyveri* species (Vakirlis et al. 2016). This suggests major differences in the regulation of crossover formation and patterning between species having and missing the ZMM pathway.

The genomic era now allows precise determination of genome-wide meiotic recombination landscapes and frequencies virtually in any species. This information is important to better understand the mechanisms controlling crossover formation, crossover patterning, and to understand how recombination landscapes and frequencies evolve, a matter poorly explored despite frequent variations between individuals, populations, sexes and taxa (Smukowski and Noor 2011; Stapley et al. 2017b). Genome-wide meiotic recombination landscapes have been determined in many organisms including yeasts species, flies, plants and mammals (Braun-Galleani et al. 2019; Comeron et al. 2012; Liu et al. 2018; 2019; Mancera et al. 2008; Ottolini et al. 2015; Qi et al. 2009; Smeds et al. 2016; Wallberg et al. 2015; Wijnker et al. 2013; Winzeler 1998). However, besides a few exceptions for *Drosophila* (Comeron et al. 2012; Samuk et al. 2020) such information is rarely available for numerous individuals from a same species or from closely related species precluding answering precisely evolutionary questions. In this context, we recently explored the recombination landscape of *Lachancea kluyveri*, a protoploid yeast species that diverged from the *Saccharomyces* genus more than 100 million years ago and we found striking differences with *S. cerevisiae* (Brion et al. 2017). These variations include a lower recombination rate (1.6 *vs* 6.0 crossover/Mb), a higher frequency of chromosomes segregating without any crossover and the absence of recombination on the entire chromosome arm containing the sex locus (Brion et al. 2017).

In the continuity of this work, here we determined the genome-wide recombination landscape of another *Lachancea* species, *Lachancea waltii*, that lacks *MLH2* and most ZMM genes except orthologs of *ZIP1* and *MER3* (Vakirlis et al. 2016). With 3.4 crossovers/Mb, we found that the *L. waltii* recombination rate is intermediate between the *S. cerevisiae* and the *L. kluyveri* species. Despite the absence of Mlh2 that restrains the extent of gene conversion tracts in *S. cerevisiae, L. waltii* gene conversion tract lengths are similar to those of *S. cerevisiae*. Remarkably, we found no conservation of the recombination hotspots across the *Lachancea* genus unlike what was observed in the *Saccharomyces* genus (Lam and Keeney 2015; Tsai et al. 2010). Finally, coherent with the loss of several ZMM proteins, the crossover interference signal in *L. waltii* is minimal and much reduced compared to *S. cerevisiae*.

## Results

### Genetic diversity across the *Lachancea waltii* species

To determine the meiotic recombination landscape of the *L. waltii* species, we first explored the genetic diversity of a collection of natural isolates in order to identify two parental polymorphic isolates to generate a diploid hybrid and its corresponding meiotic progeny. The collection used for resequencing consists of all the seven currently available isolates of *L. waltii*, including the CBS 6430 reference isolate. This strain was isolated in Japan from the tree *Ilex integra* (Kodama and Kyono, 1974) and its genome was fully sequenced, assembled and annotated previously (Kellis et al. 2004; Vakirlis et al. 2016). It consists of 10.2 Mb spreading over eight chromosomes (A to H). The six remaining *L. waltii* natural isolates come from either trees or insects from Canada (Table S1A). We resequenced the full genome of each isolate via a short-read Illumina sequencing strategy, generating a 75-fold genomic coverage on average. The reads associated with each sample were mapped against the CBS 6430 reference sequence. Variant calling allowed to detect a total of 954,071 SNPs distributed across 227,304 polymorphic sites (Table S2). In addition, the reads coverage profiles showed the absence of aneuploidy in this set of strains and a single segmental duplication of approximately 130 kb was detected on chromosome A in the CBS 6430 strain (Figure S1). Overall, we observed a genetic divergence of approximately 1.5% (or 160,000 SNPs) between the six strains isolated from Canada and the reference isolate, highlighting the presence of two subpopulations. The Canadian subpopulation shows a sequence divergence ranging from 0.31 to 0.65% (Figure S2; Table S3).

### Extensive LOH tracts in *L. waltii* hybrids

The *L. waltii* species propagates vegetatively as a haploid. Opposite mating types are seen, but not at the same frequency, diploids are rare, and sporulation is infrequent. (Di Rienzi et al. 2011). *L. waltii* mates and sporulates on the same malt-agar medium (Di Rienzi et al. 2011). We generated hybrids between the LA128 and LA136 strains as well as between the LA133 and LA136 isolates owing to their SNP density, approximately 0.64 and 0.4%, respectively. We isolated multiple hybrids from both crosses at 48h and 72h time points after the cross. We sequenced nine hybrids of the LA128 and LA136 cross (four at 48h and five at 72h) and six hybrids of the LA133 and LA136 cross (two at 48h and four at 72h).

The analysis of the genome of these hybrids highlighted a plethora of large regions of loss of heterozygosity (LOH) (Figure S3). The LA128/136 hybrids had a mean of 12 (±0.9) LOH tracts, and the LA133/136 hybrids had a mean of 21 (±2.1) LOH tracts. Because *L. waltii* mates and sporulates on the same medium, all these LOH regions could result from abortive meiosis also called “return to growth (RTG)”, or from intra-tetrad mating (Brion et al. 2017; Dayani et al. 2011; Laureau et al. 2016). RTG seems more likely since mating frequency is low. Either way, formation of LOH regions seems unavoidable so far as *L. waltii* mates and sporulates on the same medium. In addition, aneuploidies of chromosomes B and D were also prevalent in both hybrid backgrounds. Extra copies of chromosome B or D were observed five and eight times, respectively, among the nine LA128/136 hybrids. In the six LA133/136 hybrids, extra copies of chromosome B or D were observed once and four times, respectively.

### *L. waltii* recombination population

LOH tracts and aneuploidies preclude optimal detection of recombination events. However, their compulsory presence in *L. waltii* hybrids led us to choose a LA128/136 hybrid that accumulated the least number of LOH tracts (n=11) and that is aneuploid for chromosome B and D to make a segregant population and determine the *L. waltii* meiotic recombination landscape (Peltier et al. 2021) (Figure 1). The 11 LOH tracts encompass five out of the six euploid chromosomes and represent a total of 1.64 Mb. The LA128/LA136 hybrid contains 64,628 polymorphic sites distributed across the eight chromosomes. This corresponds to a divergence of ∼0.64 %, which is in the same order of magnitude as those of the hybrids used to study meiotic recombination landscapes in *S. cerevisiae* and *L. kluyveri* (Mancera et al. 2008; Chen et al. 2008; Martini et al. 2011; Brion et al. 2017). The LA128/LA136 hybrid shows ∼30% sporulation efficiency after 72 hours on malt agar medium, with 71% spore viability and 42.4% of full viable tetrads (Table S4).

**Figure 1.**
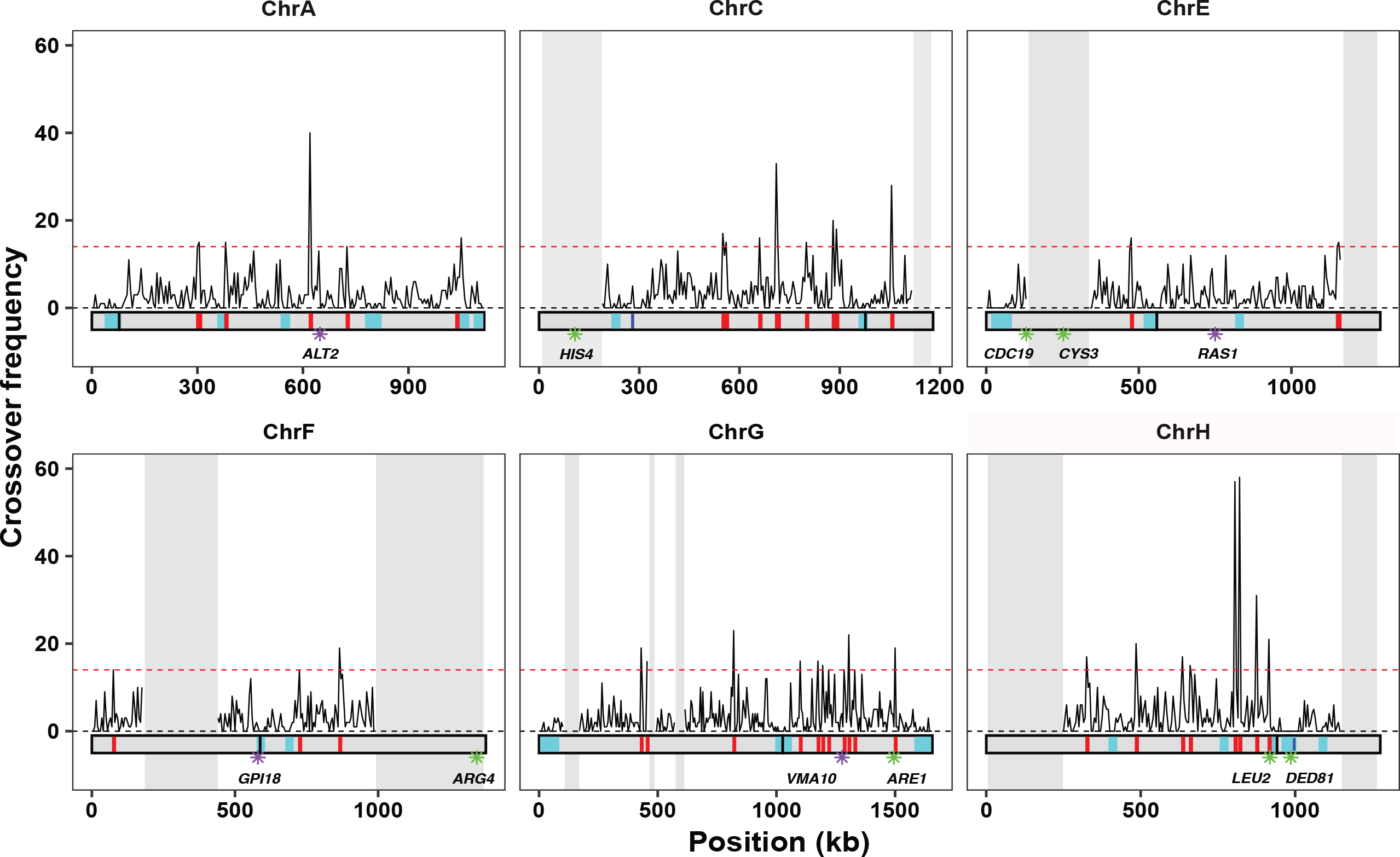
*L. waltii* recombination map. Density of crossovers along the genome using a 5 kb window. Grey shaded areas represent LOH regions from the parental hybrid. Horizontal dotted red line represents crossover hotspot significance threshold (see methods). Under the density plot, crossover coldspots and hotspots are shown in cyan and red, respectively. For each chromosome a black dash represents centromere position. Blue dashes represent MAT locus on chromosome C and rDNA locus on chromosome H. Genes of *L. kluyveri* and *S. cerevisiae* within hotspot positions according to (Mancera et al. 2008; Brion et al. 2017) are represented by purple and green star, respectively.

We generated 768 segregants from 192 full viable tetrads. We genotyped this F1-offspring population by Illumina whole genome sequencing and identified recombination events based on 50,274 SNPs parental inheritance, disregarding chromosome B and D as well as the 11 LOH regions for the analysis of recombination. Overall, we generated a recombination map for 5.7 Mb out of the 10.2 Mb of the genome. In order to validate the 50,274 SNPs used for genotyping, we computed their 2:2 segregation and their pairwise recombination fractions. This analysis revealed the presence of a reciprocal translocation between chromosomes A and F in the parental strains compared to the CBS 6430 reference genome (Figure S4A; 730 kb on chromosome A and 670 kb on chromosome F). This translocation could also be identified through genome assemblies and is shared by all the Canadian isolates (Figure S4B). Crossovers and noncrossovers detected around the translocation breakpoints on chromosomes A and F (1 kb window on chromosome A, 12.2 kb window on chromosome F empirically determined based on the range of misaligned reads around the breakpoints) were excluded from downstream analyses. Overall, the median distance between consecutive markers in euploid heterozygous regions of the LA128/136 hybrid is 159 bp. These markers are distributed rather evenly across the genome, giving a resolution comparable to previous studies in *S. cerevisiae* and *L. kluyveri* (Brion et al. 2017; Chen et al. 2008; Mancera et al. 2008; Martini et al. 2011).

### *L. waltii* recombination landscape

In the 192 four-spore viable tetrads analyzed, a total of 4,049 crossovers and 1,459 noncrossovers were identified, leading to a median of 19 crossovers and 7 noncrossovers per meiosis (Table S5). Additionally, a detectable gene conversion tract was observed in 73% of the crossovers. The crossover and noncrossover rates were 3.4 crossovers/Mb and 1.2 noncrossover/Mb, respectively. Based on this, we estimated nearly 34 crossovers and 12.5 noncrossovers per meiosis genome wide in *L waltii*. With an intercept around 1 (0.72), the average frequency of crossovers per chromosome exhibits a linear relationship with LOH corrected chromosome size (Figure 2A). By contrast to *S. cerevisiae*, crossover density has no significant inverse correlation with chromosome size in both *L. waltii* and *L. kluyveri* (Figure 2B). Similar results have been seen in crossover interference mutants in *S. cerevisiae* (Anderson et al. 2015; Chakraborty et al. 2017).

**Figure 2.**
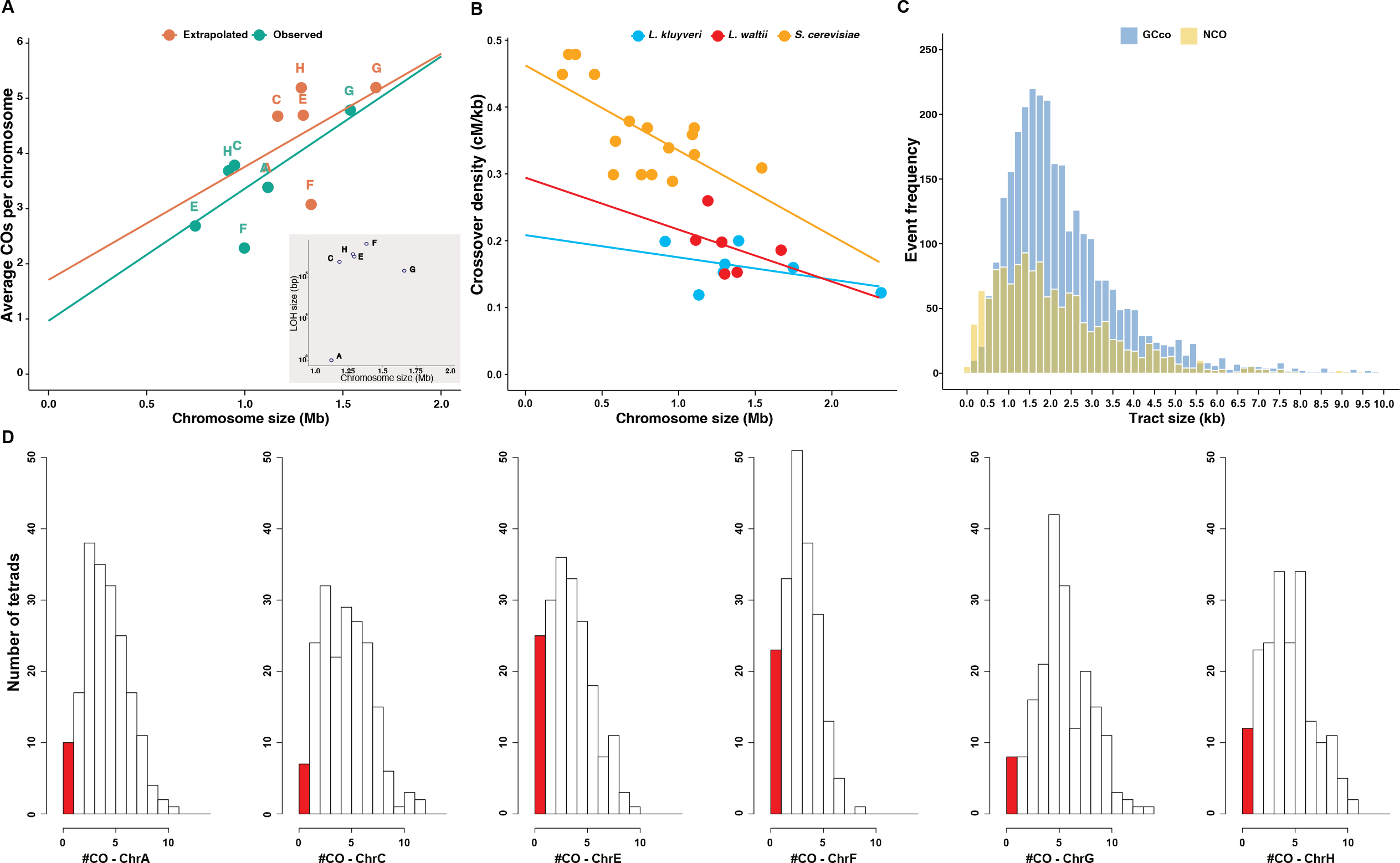
*L. waltii* recombination events. **A.** Frequency of crossovers in *L. waltii* are related to chromosome size. The ‘observed’ version corresponds to the average number of CO detected compared to to the studied fraction of each chromosome, *i*.*e*. excluding the LOH regions (intercept=0.72, slope=2.65 × 10^−6^). The ‘extrapolated’ version represents the corrected number of COs relative to the actual chromosome size (intercept=1.4, slope= 2 × 10^−6^). The corrected CO number was estimated by extending the CO rate (3.4 CO/Mb) to the whole chromosome. [Inset: cumulative sizes of regions under LOH relative to the actual chromosome size]. **B**. Recombination rate (cM/kb) plotted against chromosome size for *L. waltii* with the extrapolated values (R=-0.71, p=0.14), *L. kluyveri* (R=-0.21, p=0.66) and *S. cerevisiae* (R=-0.6, p=0.6, p= 1 × 10^−4^). *L. kluyveri* data from Brion et al. 2017, *S. cerevisiae* data pooled from Chakraborty et al. 2017 and Anderson et al. 2015. **C**. Distribution of gene conversion tract sizes associated to crossovers (GCco) and noncrossovers. **D**. Histogram representing the frequency of crossovers per meiosis in each *L. waltii* chromosome. The red bars represent instances where no crossover was detected on the chromosome (E_0_).

Most ancestral heterozygous SNPs (84.8%) segregated 2:2 in *L. waltii*. The median sizes of conversion tracts associated to crossovers (GCco) and noncrossovers were 2.0 kb and 1.8 kb, respectively, and were not significantly different (Wilcoxon test, p<0.0001; Figure 2C and Table S6). Such GCco and noncrossover tract lengths are identical to those from *S. cerevisiae* but shorter than those from *L. kluyveri* (2.8 and 3.0 kb for GCco and noncrossovers, respectively; Wilcoxon test, p<0.0001).

Approximately 3% of *S. cerevisiae* meioses exhibit chromosomes that segregate without crossover, *i*.*e*. non-exchange or E_0_ chromosomes (0 for no exchange), but only small chromosomes are impacted (Mancera et al. 2008; Chakraborty et al. 2017). In contrast, almost 45% of *L. kluyveri* meioses had an E_0_ chromosome, disregarding chromosome C where crossovers are suppressed in a large region of the left arm (Brion et al. 2017). In *L. waltii*, the incidence of E_0_ chromosomes is also high, with 22.9% of meioses displaying at least one and 8.3% exhibiting two or more E_0_ chromosomes (Figure 2D). This result comes with the caveat that LOH in the parental hybrid may exaggerate the E_0_ estimates. However, *L. waltii* 1.12 Mb long chromosome A, which displayed no LOH events, was found to be E_0_ in 5.2% of the meioses. This contrasts with *S. cerevisiae* where chromosomes of equivalent size never show zero crossover (Mancera et al. 2008; Krishnaprasad et al. 2015; Chakraborty et al. 2017; Zhang et al. 2014a). Overall, unlike *S. cerevisiae*, both *Lachancea* species seem to be able to segregate accurately a high frequency of large chromosomes without crossover. Interestingly, the frequency of E_0_ chromosomes in *L. waltii* was lower than in *L. kluyveri*, despite lacking several pro-crossover ZMM genes.

### *L. waltii* recombination hotspots and coldspots

We computed the local recombination rate using the 4,049 single crossovers in order to identify potential crossover hotspots or coldspots (Figure 1). To compare with *L. kluyveri*, recombination hotspots were identified using the same window size of 5 kb and significance assessed using permutated datasets (10^5^ permutations). Recombination coldspots were identified with the same approach using a 20 kb window size to increase detection power. We identified 21 coldspots and 37 hotspots of recombination.

As observed in many species including *S. cerevisiae* and *L. kluyveri*, recombination coldspots include the eight centromeres (Mancera et al. 2008; Brion et al. 2017; Braun-Galleani et al. 2019; Liu, Maclean, and Zhang 2019), the surrounding of the rDNA locus (Gottlieb and Esposito 1989; Vader et al. 2011) and all LOH-free sub-telomeric regions (chromosomes A, E, G), except the chromosome F left sub-telomere (Liu, Maclean, and Zhang 2019; Pan et al. 2011). In addition, nine recombination coldspots are located away from any centromere, subtelomere and the rDNA locus. Interestingly, in five of the nine recombination coldspots, we identified four genes which *S. cerevisiae* orthologs are connected to DNA metabolism. LAWA0A06634g and LAWA0E09516g are the orthologs of the *DMC1* and *MEI5* genes, which encode the meiotic specific recombinase and one of its accessory factors, respectively. LAWA0F07800g is the ortholog of *SLX4*, which encodes a scaffold protein that controls and coordinates the action of multiple structure specific endonucleases and interacts with other DNA repair factors (Guervilly and Gaillard, 2018)). Finally, LAWA0H08790g is the ortholog of *IRC8* which deletion is associated with an increased level of Rad52-GFP foci potentially revealing a role in DNA metabolism. This result is in accordance with previous evidence in *S. cerevisiae* meiotic DSB map analysis, which showed that meiotic genes tend to be protected against meiotic DSBs (Pan et al. 2011).

A striking feature of *L. kluyveri* recombination landscape was the absence of recombination on the entire left arm of the chromosome C (Lakl0C-left) (Brion et al. 2017). The synteny of part of this large 1-MB region is conserved over 600 kb between *L. kluyveri* and *L. waltii*. By contrast to *L. kluyveri*, the *L. waltii* region orthologous to the Lakl0C-left exhibits a recombination rate (3.56 crossovers/Mb) similar the rest of the genome.

Finally, we identified about five times more crossover hotspots in *L. waltii* compared to *L. kluyveri* (37 *vs* 7, respectively) using the same method and threshold. This likely comes from a *L. waltii* dataset with a higher detection power due to four times more meioses analyzed and an about two-fold higher recombination rate. Among the seven *L. kluyveri* crossover hotspots, *PIS1* and *PRM2* do not have annotated orthologs in *L. waltii* and *OST6* ortholog is located on *L. waltii* chromosome D which was discarded from our analysis because of the presence of an aneuploidy. None of the four remaining *L. kluyveri* crossover hotspots are crossover hotspot in *L. waltii*. Comparison of *L. waltii* crossover hotspots with *S. cerevisiae* ones (Gerton et al. 2000; Mancera et al. 2008) revealed that only the *ARE1* associated crossover hotspot is conserved. In conclusion, crossover hotspots are overall not conserved between *L. waltii* and *L. kluyveri* or *S. cerevisiae*.

### *L. waltii* crossovers exhibit interference

Crossover interference is an interactive process between crossovers along chromosomes, enabling non-random distribution of crossovers (Zickler and Kleckner, 2015). We analyzed crossover interference genome-wide using the coefficient of coincidence (CoC) method (Materials and Methods and see Anderson et al. 2015). CoC is the ratio of the observed frequency of crossovers in two consecutive chromosomal intervals out of the expected frequency of such double crossovers if they arose independently in the two intervals. In *L. waltii*, crossover interference (1-CoC) is only positive for the smallest interval size of 25 kb, while in *S. cerevisiae* it is positive up to a 100 kb interval size (Figure 3A). Only chromosomes A, E, and H display interference over a 25-kb interval, but chromosomes C, F, and G do not (Figure 3B). Unfortunately, the available *L. kluyveri* crossover dataset did not allow such a CoC analysis to be performed due to the small sample size.

**Figure 3.**
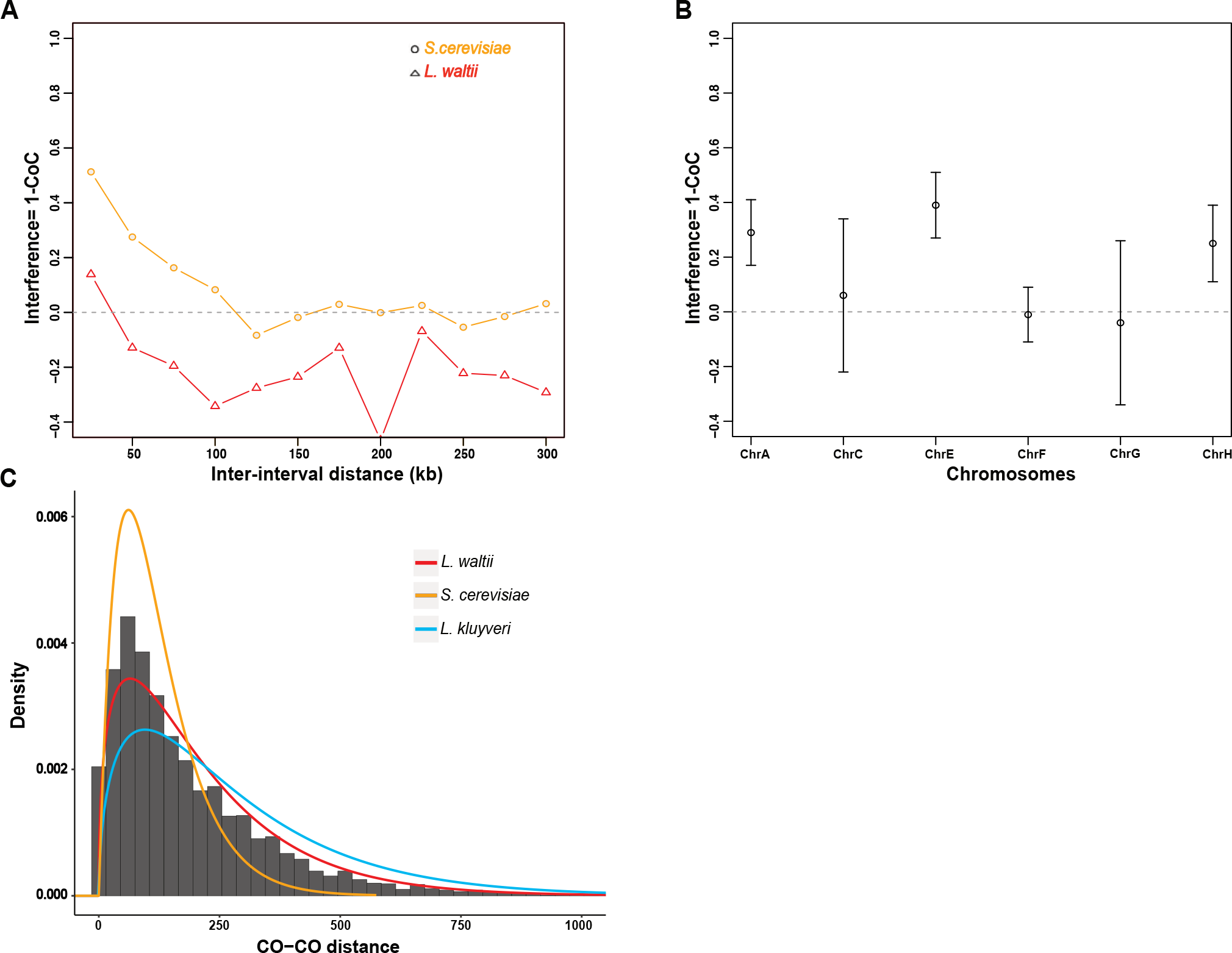
Crossover interference in *L. waltii*. **A**. Crossover interference (1-CoC) calculated for a bin size of 25 kb and inter-interval distance of 25 kb (i.e., for adjacent intervals) in *L. waltii* (triangle) and *S. cerevisiae* (circle). **B**. Chromosome-wide interference (1-CoC) in 192 *L. waltii* tetrads calculated for a bin size of 1/ (5*(mean inter crossover distance)) for adjacent bins using MAD patterns (White et al. 2017). **C**. Distribution of distances between pairs of adjacent crossovers in *L. waltii* (dark grey). A gamma law fit of this distribution is represented, as well as a gamma law curve for *S. cerevisiae* and *L. kluyveri* using parameters described in Anderson et al. 2011 and Brion et al. 2017, respectively. Gamma parameters were as follows: for *L. waltii*, k=1.23 and ⍬=145.9, for *S. cerevisiae*, k=1.96, ⍬=61.7 and for *L. kluyveri*, k=1.47, ⍬=187.

The strength of interference can also be determined by modelling inter-crossover distances using a gamma distribution: γ = 1 indicates no interference, whereas γ > 1 corresponds to positive interference. The γ-values of the gamma-fit distributions for *S. cerevisiae, L. kluyveri* and *L. waltii* are 1.96, 1.47 and 1.23, respectively (Figure 3C) (Brion et al 2017). The gamma-fit distribution of inter-crossover distances in *L. waltii* is significantly different from both of *S. cerevisiae* and *L. kluyveri* as well as from a gamma law of shape=1 (K-S test, p<0.0005). Overall, the CoC method and the modelling with a gamma distribution support the presence of crossover interference in *L. waltii* with a reduced strength relative to *S. cerevisiae*.

We further modelled the crossover distribution in *L. waltii* with the CODA package, using the two-pathway model that incorporates contributions from both interfering and non-interfering crossover pathways (Gauthier et al. 2011). This analysis suggests that only 7.7% of the crossovers in *L. waltii* are non-interfering, similar to *S. cerevisiae* with 5.8% (Table S7). Altogether, our data suggests that most crossovers in *L. waltii* interfere, but interference barely extends beyond 25 kb.

## Discussion

### Mechanism of LOH formation: an under-estimate of the importance of RTG in every yeast species?

Despite our efforts to reduce the time cells spent on malt agar, the medium that allows both mating and sporulation of *L. waltii*, we could never recover hybrids without any LOH. The two most likely scenarios for the formation of LOH regions in the parental hybrids are RTG and sporulation induction followed by inter-spore mating. A leaky activity of Spo11 before meiotic entry is also plausible. Interestingly, *L. kluyveri* also showed a related phenotype that we attributed to RTG, but for which inter-spore mating could not formally be ruled out (Brion et al. 2017). Further experiments are necessary to distinguish between these two routes in both species. In the case of inter-spore mating, it would suggest that at least two rounds of meiosis occur in a row systematically in *L. waltii*. In the case of RTG, it would suggest that there is frequent reversion of meiotic entry in *L. waltii*. In the latter case, the evolutionary importance of RTG should be revisited as an efficient and frequent way to shuffle parental genomes without forming gametes (Laureau et al. 2016; Mozzachiodi et al. 2021).

### Which nuclease(s) make(s) crossovers in *L. waltii*?

So far, interfering crossovers were known to result from the ZMM pathway which Holliday junction resolvase activity comes from the action of the Mlh1-Mlh3 endonuclease optimized by the presence of Exo1 (independently of its nuclease activity), Msh4-Msh5, RFC and PCNA (Rogacheva et al. 2014; Kulkarni et al. 2020; Cannavo and Cejka 2020). Interfering crossovers are also known to form in the context of the synaptonemal complex. Zip2, Zip4 and Spo16 that form the ZZS complex promote interfering crossovers and connect both the chromosome axis and Holliday junctions (HJs) with high affinity, as well as the other interfering crossover promoting factor Zip3. The absence of the ZZS complex, Zip3 and Msh4-5 in *L. waltii* questions the involvement of Mlh1-3 in the formation of interfering crossovers. Either Mlh1-3 evolved to resolve HJ junctions in the absence of its accessory factors or this activity has been taken over by other structure specific nucleases. Noteworthy, the scaffold protein Slx4 connecting several structure specific nucleases lies in a crossover coldspot in *L. waltii*. This feature is puzzling since recombination hotspots have been shown to be associated with purifying selection in *S. cerevisiae*, which by inference suggests that Slx4 is not under purifying selection or at least less than other regions of the genome (Weber and Hurst, 2009). Whether or not this underscores a role of Slx4 in the resolution of meiotic recombination intermediates and hence crossover formation in *L. waltii* remains to be determined.

In addition to the ZMM proteins, Mlh2 is also absent from *L. waltii*. In combination with Mlh1, it forms the MutLß complex that limits D-loop extension and likely the subsequent gene conversion tracts associated to both noncrossovers and crossovers in *S. cerevisiae* (Duroc et al. 2017; Vernekar et al. 2021). Interestingly, we found that the gene conversion tract lengths are of similar sizes in *L. waltii* and *S. cerevisiae*, and smaller than in *L. kluyveri*. This suggests that the MutLß role in limiting D-loop extension is dispensable in *L. waltii* to generate short conversion tracts, and hence that subtle differences exist in the biology of recombination intermediates between *L. waltii, L. kluyveri* and *S. cerevisiae*.

### E_0_ chromosomes

The high fraction of E_0_ chromosomes in *L. waltii* contrasts with the low level observed in *S. cerevisiae*. This could come from the lack of recombination information from a large fraction of the genome corresponding to LOH regions in the staring hybrid. However, this argument can be discarded by chromosome A (and G to a lesser extent) that has no LOH and that still shows about 5% E_0_ among the 192 tetrads. This high fraction of E_0_ is similar to that observed in *L. kluyveri* chromosomes except for the mating type chromosome where the fraction of E_0_ reached 50% likely because of the absence of recombination on its 1 Mb-long left arm. This supports the existence of an efficient distributive chromosome segregation mechanism in *L. waltii*, which would be more important than in *S. cerevisiae*, as postulated for *L. kluyveri* (Brion et al. 2017).

### Crossover interference in *L. waltii*

Although relatively weak compared to other species such as *C. elegans*, crossover interference exists in *S. cerevisiae* and has been shown to rely on Top2 and Red1 through sumoylation (Zhang et al. 2014). Top2 and Red1 are conserved in *L. waltii*. However, Zip2, Zip3, Zip4, Spo16, Msh4 and Msh5, that likely play a role in crossover interference implementation as the corresponding mutants accumulate non-interfering crossovers in different species (Pyatnitskaya et al. 2019), have been lost in *L. waltii*. As a consequence, it was expected that *L. waltii* was defective for crossover interference. Interestingly, we found evidence of crossover interference in *L. waltii*, albeit with reduced strength compared to *S. cerevisiae*. Although the optimal metrics for crossover interference measure is meiotic chromosome axis length, crossover interference hardly extends beyond 25 kb in *L. waltii*. This size is in the range of size of a single chromatin loop in *S. cerevisiae* and *L. kluyveri* (Muller et al., 2018; Schalbetter et al., 2019). One explanation for this reduced but not null crossover interference could be that crossover interference is normally established but not properly implemented due to the absence of Zip2, Zip3, Zip4, Spo16, Msh4 and Msh5. Alternatively, it is possible that the crossover interference signal detected in *L. waltii* uniquely results from the non-uniform distribution of meiotic DSBs. Indeed, previous studies already reported weak but significant interference between crossover precursors both in tomato and in mouse (Anderson et al. 2001; Berchowitz and Copenhaver 2010; de Boer et al. 2006). In addition, a modeling study also supported an even patterning of meiotic DSBs in *S. cerevisiae*, and further showed that even DSB patterning *per se* was enough to generate a positive CoC signal (Zhang et al. 2014). Overall, the crossover interference signal detected in *L. waltii* is compatible with a signal solely resulting from an even distribution of meiotic DSBs. It supports the already formulated view that crossover interference is multilayered, with a short-range component influenced by positive interference between DSBs and at least a long-range component (Berchowitz and Copenhaver, 2010). Positive DSB interference has been shown to rely on Tel1 in *S. cerevisiae* (Anderson et al. 2015; Garcia et al. 2015). The *L. waltii* context supports that a long-range component controlling crossover interference involves some or all Zip2, Zip3, Zip4, Spo16, Msh4 and Msh5 in *S. cerevisiae*. Another layer of regulation exists at least in *A. thaliana* and *C. elegans* where the central element of the synaptonemal complex was shown to play a role in establishing strong crossover interference (Libuda et al. 2013; Capilla-Pérez et al. 2021; France et al. 2021).

### Crossover hotspots conservation

DSB hotspots, that determine the location of recombination hotspots in many species including *S. cerevisiae* and likely any budding yeast, and recombination hotspots, are relatively stable across *Saccharomyces* species (Lam and Keeney 2015; Tsai et al. 2010). Conservation of recombination hotspots has also been observed across flinch species, where DSB hotspots are also determined simply by nucleosome free regions and where PRDM9 is absent (Singhal et al. 2015). Such a situation contrasts with the lack of recombination hotspots conservation we observed here between *L. kluyveri* and *L. waltii*, two *Lachancea* species that diverged about 80 million years ago. A possible explanation is that the genomes of the *Saccharomyces* yeasts analyzed are mostly colinear, with very limited genomic rearrangements, while about 130 chromosome translocations and inversions have been inferred between the extent *L. kluyveri* and *L. waltii* species (Vakirlis et al. 2016). Although we postulated a similar argument for the lack of conservation of recombination hotspot between *L. kluyveri* and *S. cerevisiae*, the present study illustrates that the loss of recombination hotspots can occur at a much-reduced evolutionary time such as the one between *L. kluyveri* and *L. waltii*.

## Materials and methods

### Yeast strains and growth conditions

The *Lachancea waltii* natural isolates and the modified strains used in this study are described in Table S1A-B. Components for cell culture medium were purchased from MP biomedicals. Strains were grown in standard YPD medium supplemented with G418 (200 μg/ml) or nourseothricin (100 μg/ml) and agar 20 g/l for solid medium at 30°C. Mating and sporulation were performed on DYM medium (yeast extract 0.3 g/l, malt extract 0.3 g/l, peptones 0.5 g/l, dextrose 1 g/l and agar 20 g/l) at 22°C.

### Generation of parental strains

Stable haploid parental strains were obtained by replacing the *HIS3* locus with G418 or nourseothricin resistance markers. The *his3Δ::KanMX* cassette was amplified from the strain 78 using primer pairs HIS3-F/HIS3-R. The *his3Δ::NatMX* locus was amplified by a two-steps fusion PCR: the flanking regions of the *HIS3* locus were amplified from the strain LA128 using HIS3-F/HIS3-fusionTermF and HIS3-R/HIS3-fusionProm-R while the *NatMX* cassette was amplified from the plasmid pAG36 using HIS3Fusion TEFProm-F/HIS3Fusion TERterm-R. pAG36 was a gift from John McCusker (Addgene plasmid# 35126) (Goldstein and McCusker 1999).

The transformation of parental strains with the *NatMX* or *KanMX* cassettes was performed by electroporation as described by (Di Rienzi et al. 2011) using 1 μg of DNA and electroporation at 1.5 kV, 25 mF and 200 Ω using a GenePulser (Biorad). To confirm successful replacement of the HIS3 locus, colony PCR were performed using primer pairs HIS3-F/Kan-R or HIS3-F/Nat-R. All primers used are detailed in Table S8.

### Mating, sporulation, and spore isolation

For mating, LA128 (*his3Δ::KanMX)* and LA136 (*his3Δ::NatMX)* were mixed on DYM plates for 72 h at 22°C. Double resistant cells to G418 and nourseothricin were selected on YPD-G418-Nourseothericin plates. Single colonies were purified by streaking on YPD-G418-Nourseothericin plates. The diploidy in hybrids LA128 (*his3Δ::KanMX*) / LA136 (*his3Δ::NatMX*) was verified by flow cytometry.

After ploidy validation, one hybrid was selected and sporulated for 2-3 days on DYM plates at 22°C. Tetrads dissections were performed using the SporePlay (Singer Instrument) without any pre-treatment. Dissection of about 1,000 tetrads showed 2:2 segregation of the *KanMX* and *NatMX* markers.

### Genomic DNA extraction

Total genomic DNA of the *L. waltii* isolates as well as the generated hybrids were extracted using a modified MasterPure Yeast DNA purification protocol (Lucigen).

Total genomic DNA of the 768 segregants was extracted using the 96-well E-Z 96 Tissue DNA kit (Omega) following a modified bacterial protocol. Cells were grown overnight at 30°C with agitation at 200 rpm in 1 ml of YPD in 2 ml 96 deep square well plates, sealed with Breath-Easy gas-permeable membranes (Sigma-Aldrich). Cells were centrifuged for 5 min at 3,700 rpm and the cell wall was digested for 2 h at 37 °C in 800 ml of buffer Y1 (182.2 g of sorbitol, 200 ml of EDTA 0.5 M pH 8, 1 ml of beta-mercaptoethanol in a total of 1 l of H_2_O) containing 0.5 mg of Zymolase 20T. The cells were pelleted, resuspended in 225 ml of TL buffer containing OB protease and incubated overnight at 56 °C. Subsequently, DNA extraction was continued following the manufacturers protocol.

DNA concentration was measured using the Qubit dsDNA HS assay (ThermoFischer) and the fluorescent plate-reader TECAN Infinite Pro200 and DNA quality was evaluated using a NanoDrop 1000 Spectrophotometer (ThermoFischer).

### Illumina high-throughput sequencing

For the seven *L. waltii* isolates, genomic Illumina sequencing libraries were prepared with a mean insert size of 280 bp and subjected to paired-end sequencing (2×100 bp) on Illumina HiSeq 2500 sequencers by the BGI.

For the hybrids and the 768 segregants, DNA libraries were prepared from 5 ng of total genomic DNA using the NEBNext Ultra II FS DNA Library kit for Illumina (New England Biolabs). All volumes specified in the manufacturer’s protocol were divided by four. The adaptor-ligated DNA fragments of about 300-bp were amplified with 8 cycles of PCR using indexed primers. A combination of 48 i7 oligos (NEBNext Multiplex Oligos for Illumina, NEB) and 24 i5 oligos (Microsynth) were designed enabling multiplexing up to 1152-samples. After quality check using a Bioanalyzer 2100 (Agilent Technologies) and quantification using the Qubit dsDNA HS assay, 4 nM of each of the 782 libraries were pooled and run on a NextSeq 500 sequencer with paired-end 75 bp reads by the EMBL’s Genomics Core Facility (Heidelberg, Germany).

### Mapping and Single Nucleotide Polymorphisms (SNPs) calling

Sequencing reads were mapped to the *L. waltii* reference genome (obtained from the GRYC website (http://gryc.inra.fr/index.php?page=download) using bwa mem (v0.7.17). Resulting bam files were sorted and indexed using SAMtools (v1.9). Duplicated reads were marked, and sample names were assigned using Picard (v2.18.14). GATK (v3.7.0) was used to realign remaining reads. Candidate variants were then called using GATK UnifiedGenotyper.

### Segregation analysis

After variant calling, SNPs called in the LA128 and LA136 parents were first filtered (bcftools view, v1.9) to define a set of confident markers, corresponding to positions with a single alternate allele, supported by at least 10 sequencing reads in each parent and with >90% of the sequencing reads covering either the reference or alternate allele. For each segregant resulting from the LA128 and LA136 cross, SNPs located at marker positions were extracted, and parental origin was assigned based on SNP correspondence between parents and spores at those positions.

In order to validate these SNPs as markers to generate the recombination map, their segregation among the progeny was investigated. If most of the markers (67 %) follow the expected 2:2 mendelian segregation, a significant amount displays other patterns. Indeed, 20 % of the markers show 0:4 / 4:0 segregation illustrating loss of heterozygosity (LOH) events in the LA128/136 hybrid. Distribution of these 0:4 / 4:0 SNPs along the genome shows that at least 1 LOH events occurred encompassing 6 of the 8 chromosomes. In total LOH events represent 1.965 Mb of the 10.2 Mb total genome size. These LOH segments cannot be used for recombination studies and therefore have been discarded for the following analysis.

Another significant amount group of SNPs encompassing all chromosomes B and D are deviating from 2:2 segregation with a more complex pattern. By looking at the coverage along the genome in LA128/136 parental hybrid, an aneuploidy with a supplemental copy of chromosome B and D was identified, explaining deviation from 2:2 segregation. Therefore, this aneuploidy was inherited in some of the segregants showing heterozygosity for theses chromosomes. In order to keep only euploid information for recombination analysis, we did not consider chromosomes B and D in our analysis. In total, the combination of chromosomes B and D and LOH regions represents 4.5 Mb that were not considered for the recombination analysis that eventually encompassed 5.7 Mb.

### Whole genome assemblies

We used Abyss (v2.0.2) (Simpson et al. 2009) with the option ‘-k 80’ to produce de novo assemblies for all the natural isolates. For each assembly, the scaffolds corresponding to the regions of interest (chromosome A from position 725,000 to 735,000 and chromosome F from 670,000 to 690,000) were detected through blastn similarity searches and compared to the reference genome with nucmer and mummerplot (Delcher et al. 2002).

### Identification of recombination events

For each tetrad resulting from the cross between the LA128 and LA136 isolates, SNPs located at aforementioned marker positions were extracted, and parental origin was assigned based on SNP correspondence between parents and spores at those positions. The result was formatted as a segfile and used as input of the CrossOver python script (Recombine suite, Anderson et al. 2011), using default parameters and adapted chromosome sizes/coordinates to fit the reference genome of *L. waltii*.

A large amount of LOH regions were present in the parental hybrid and was passed to the studied tetrads, introducing noise in the results obtained with CrossOver. In addition, chromosomes B and D, which are prone to aneuploidies, could not reliably be analyzed for the presence of crossovers and noncrossovers. Finally, a translocation event between chromosome A and chromosome F also introduced false positive crossing-over calls at the translocation breakpoints. Thus, crossover and noncrossover events reported in chromosomes B and D (approx. 2.35 Mb), in LOH tracts (mean size: 2.17 Mb per tetrad) and at the chromosomes A-F translocation breakpoints (13.2 kb) were masked. The remaining studied genome size after masking was of 5.7 Mb.

### Determination of recombination hotspots and coldspots

The density of crossovers per interval of a set size was simulated using 10^5^ random simulations of the distribution of 4,049 crossovers, following a binomial probability model with equal chances for each event to fall into an interval. For each of the 10^5^ simulations, the maximum or minimum number of crossovers per interval was extracted and the 2000th highest or lowest value set as the threshold to define hotspots and coldspots, respectively, corresponding to p<0.02. An interval size of 5 kb was used to determine that hotspot threshold, while an interval size of 20 kb was used to increase detection power when determining the threshold for coldspots.

### Evaluation of crossover interference

A gamma law was fitted to the distribution of all inter-crossover distances in the 192 tetrads using the fitdistr R function (R, MASS package). Then the shape (k) and scale (θ) parameters of the fitted function were extracted. Crossover interference was evaluated by running a Kolmogorov-Smirnov test (using the ks.test function in R) between the distribution of all inter-crossover distances in the 192 tetrads and a gamma law of shape 1 (*i*.*e*. without interference) and scale similar to the one previously determined.

CODA (V1.2; Gauthier et al. 2011) was used to assess interference and distinguish class I crossovers (interference dependent) from class II crossovers (interference independent). Relative positions of crossover events across each chromosome were computed and used as input for CODA. The gamma-sprinkling model was used to identify the two classes, using the “two-pathway” option with default min, max and precision parameters. The projected score was selected as the score type to fit the model, with 10^6^ simulations as suggested in CODA’s help document. The hill-climbing algorithm was used as the algorithm of choice for determining optimum parameters.

## Data availability

Sequence data deposited in the European Nucleotide Archive under the accession number PRJEB39767.

## Supplementary figures

**Figure S1.**
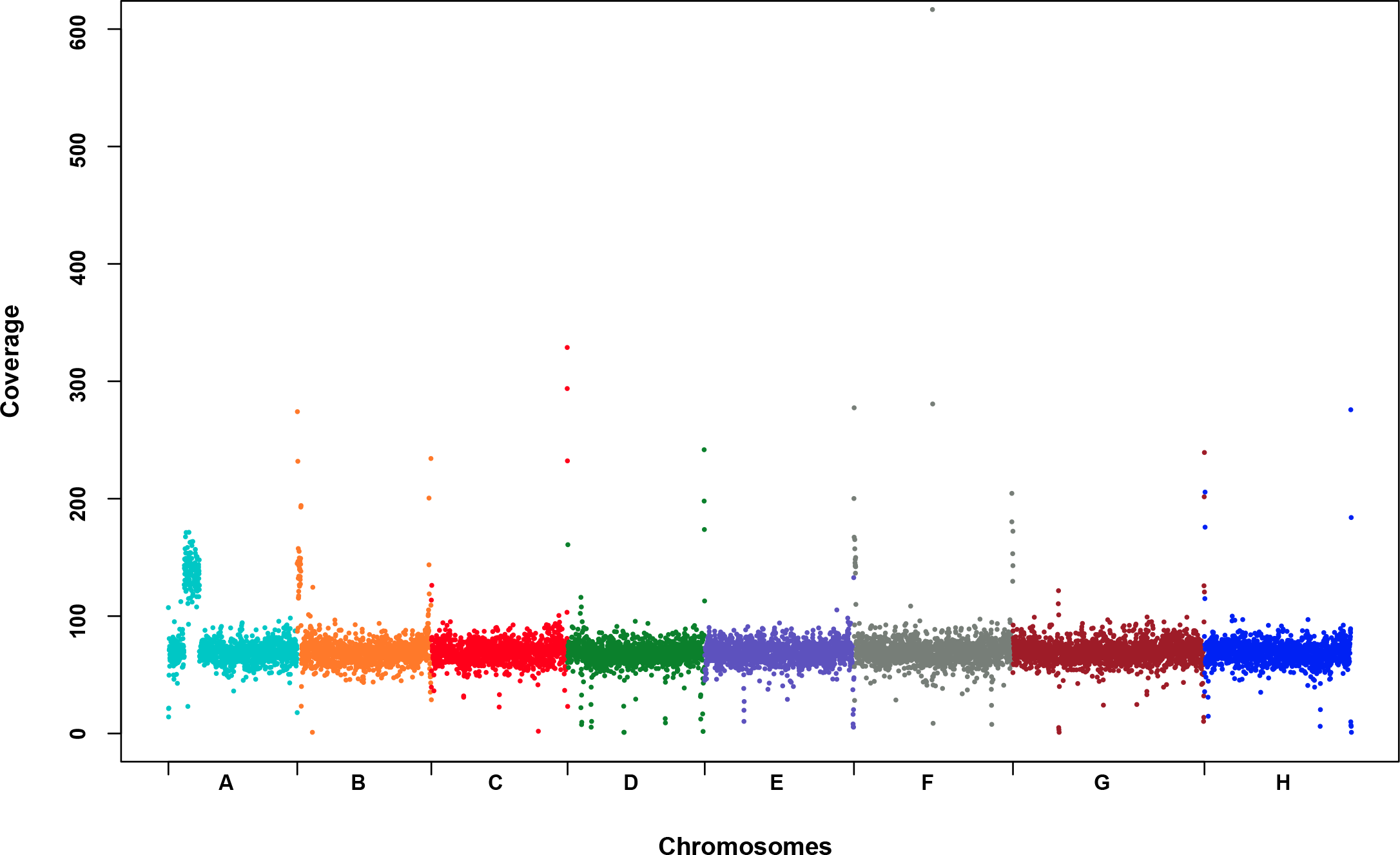
Sequencing coverage across the 8 chromosomes of the CBS6430 (LA126) reference isolate. A large 130 kb segmental duplication is detected at the beginning of the chromosome A.

**Figure S2.**
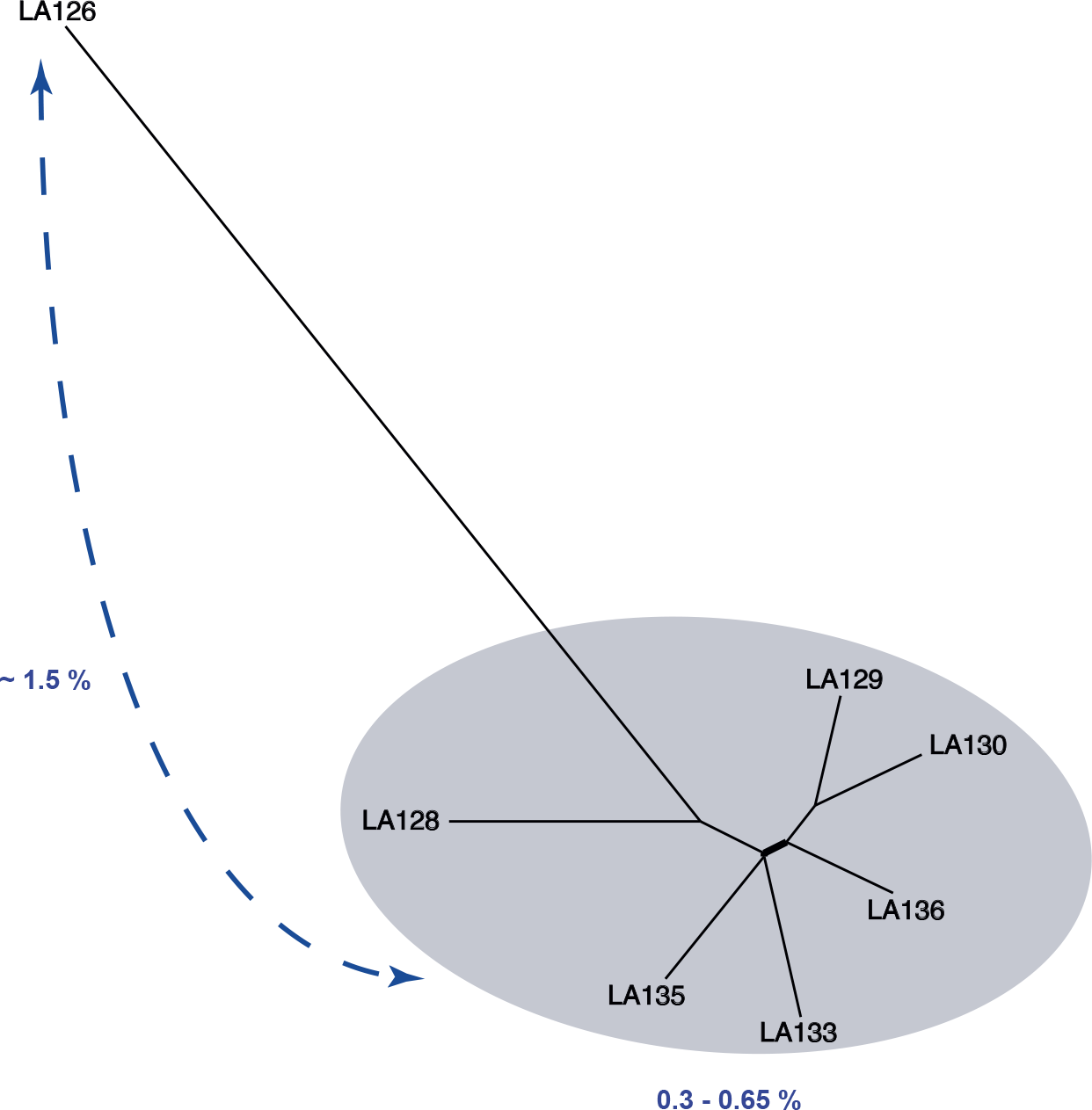
Neighbour joining tree of the studied isolates, based on 227,304 polymorphic positions. This tree underscores the presence of two main subpopulations in *L. waltii*, with LA126 lying away from the six other isolates. The relevant percentages of sequence variation between genomes are indicated in blue.

**Figure S3.**
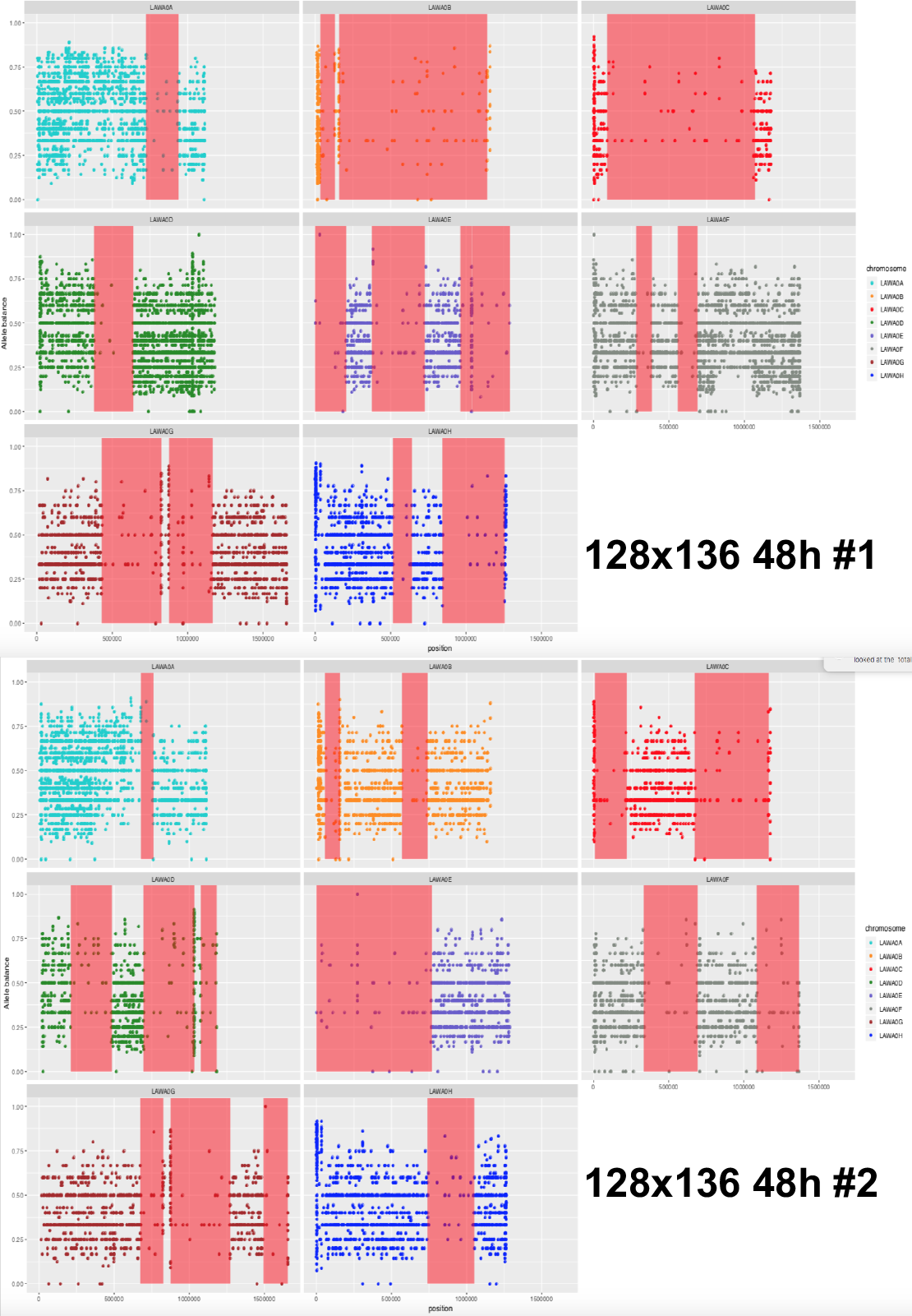
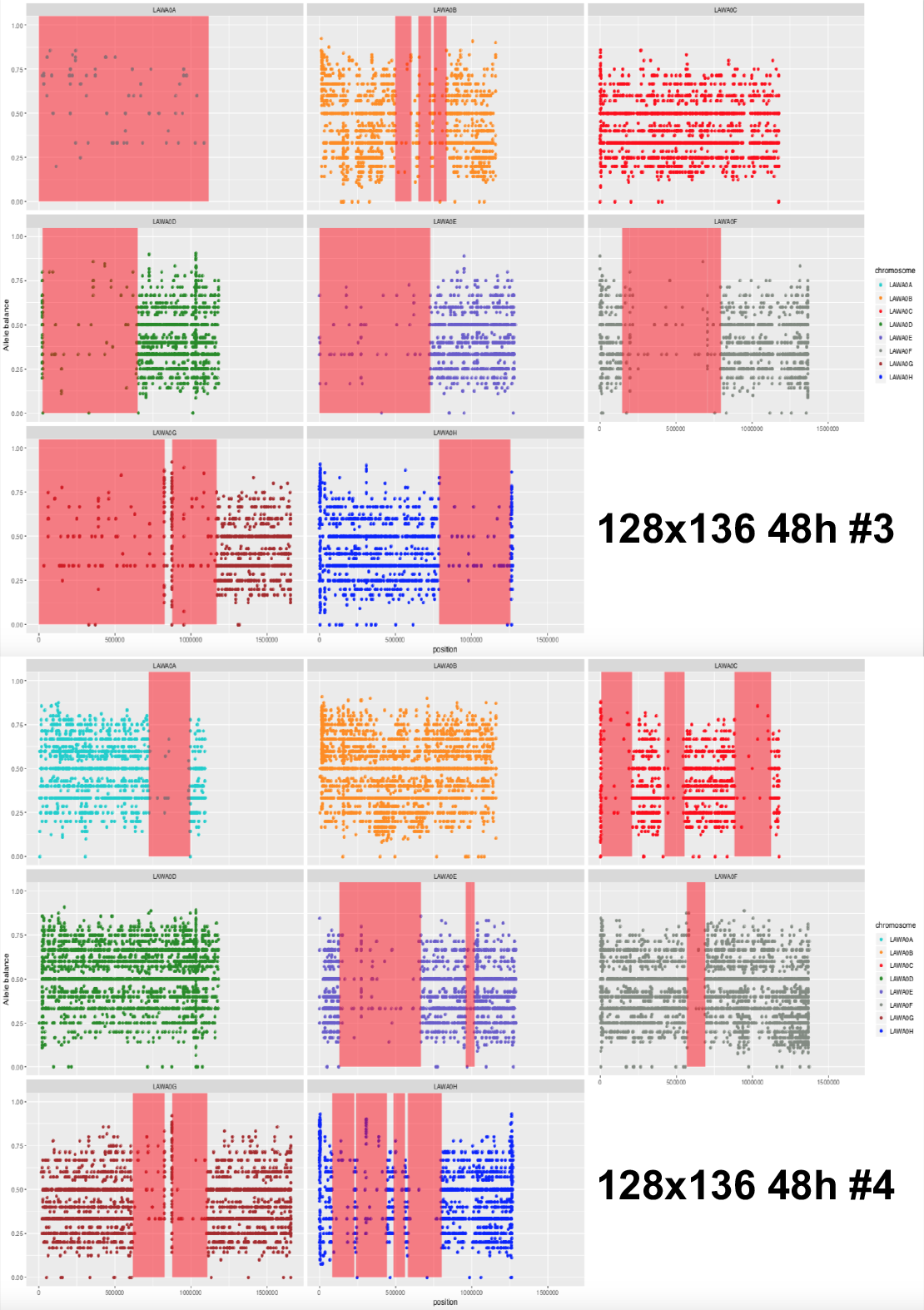
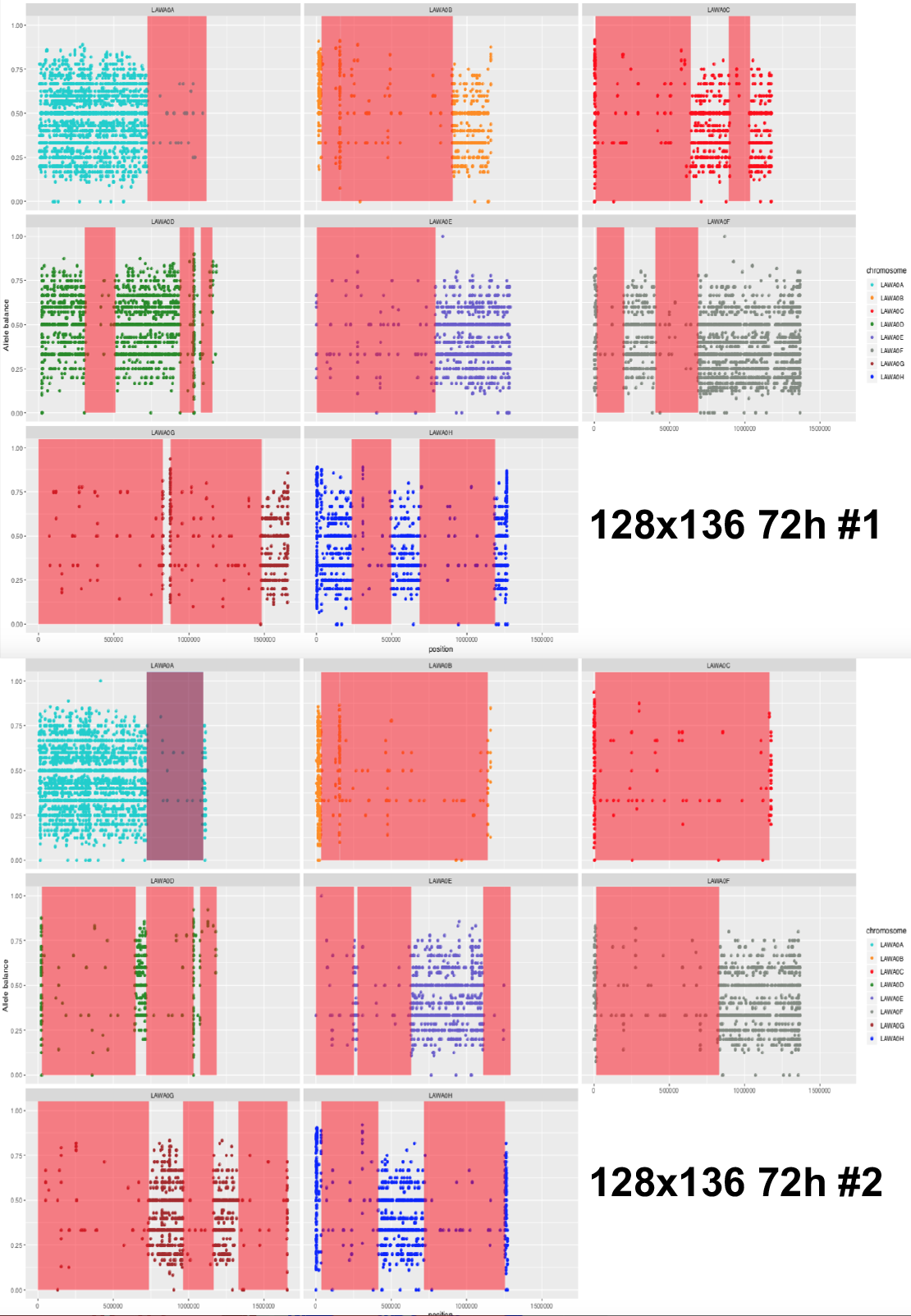
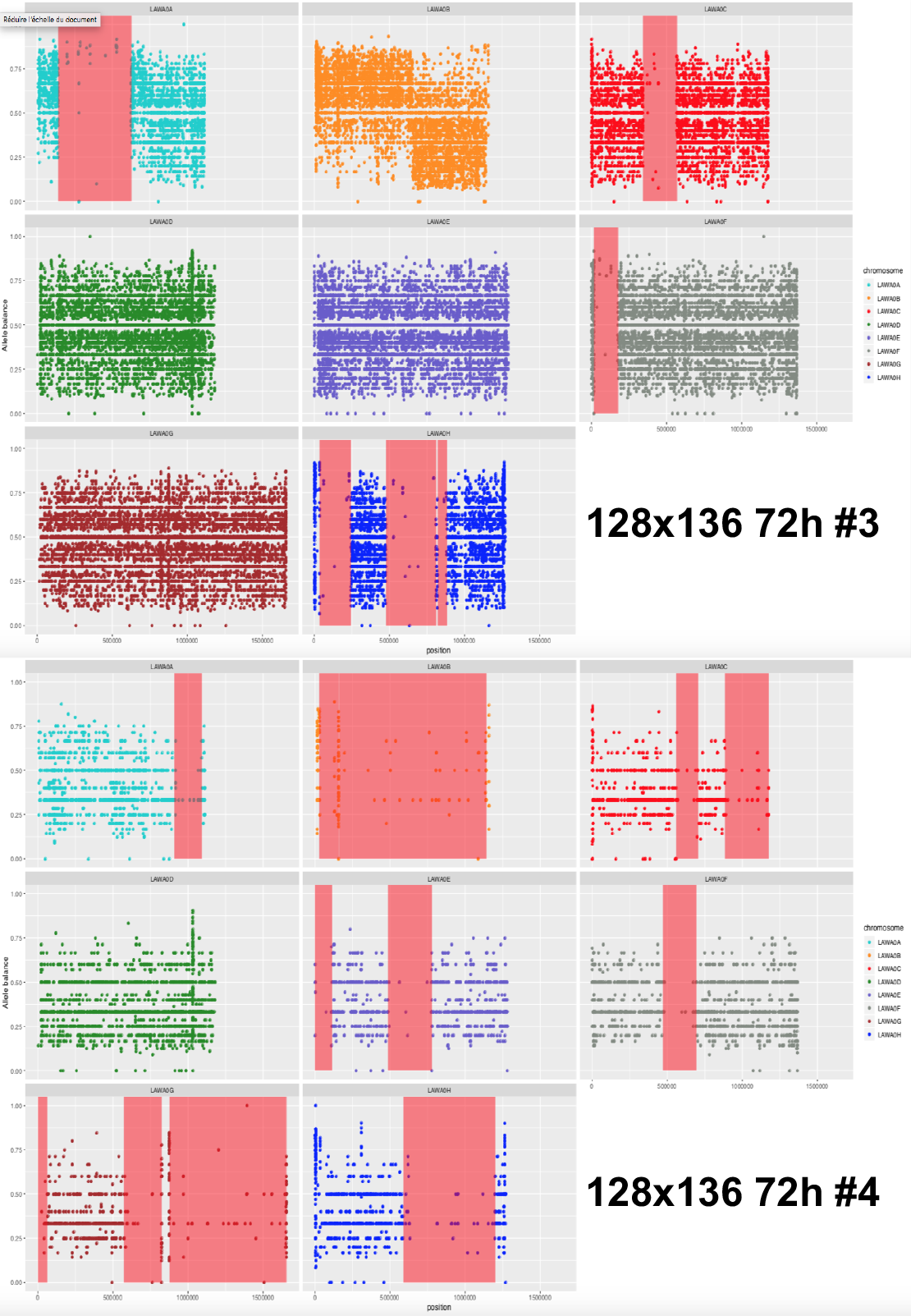
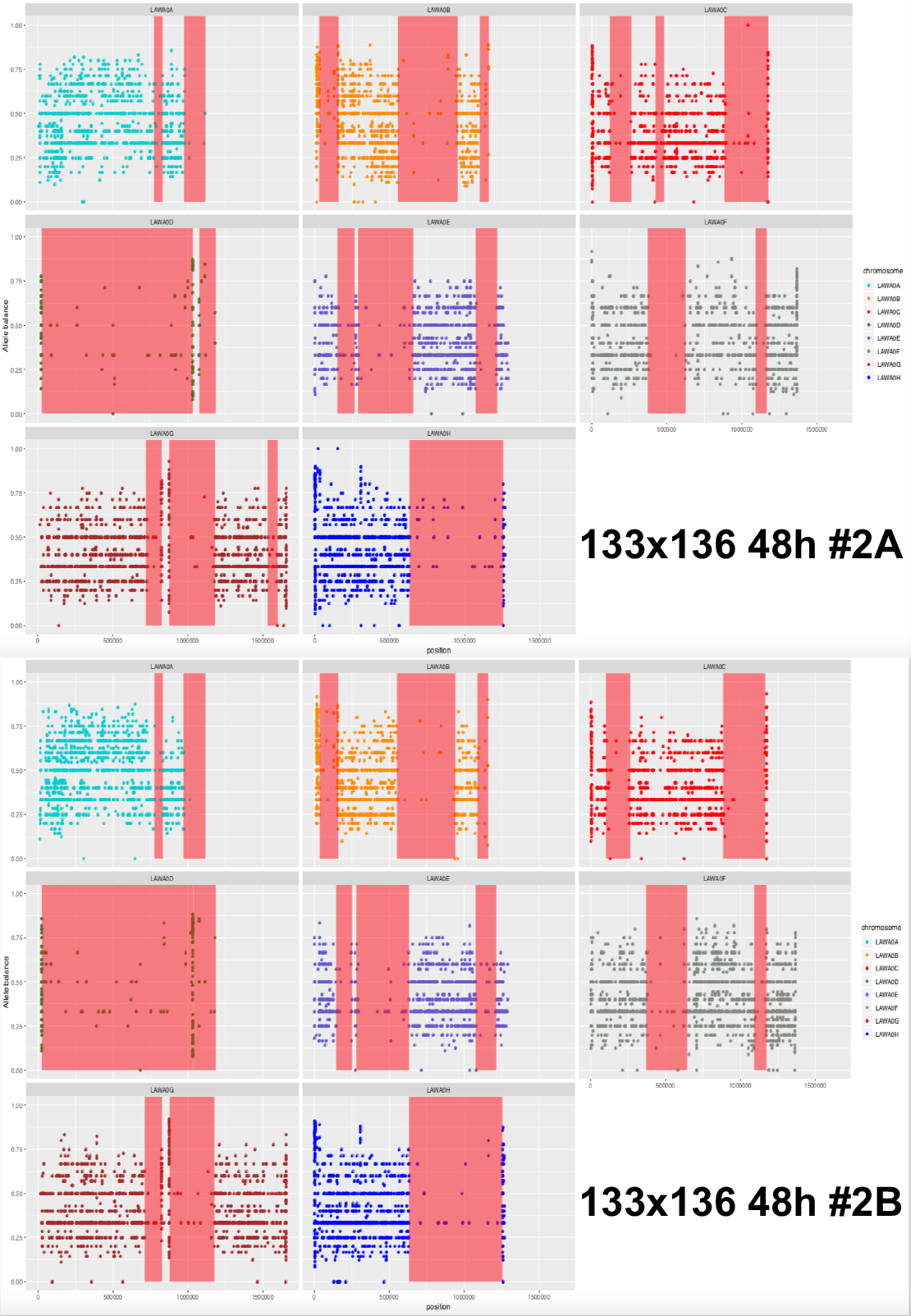
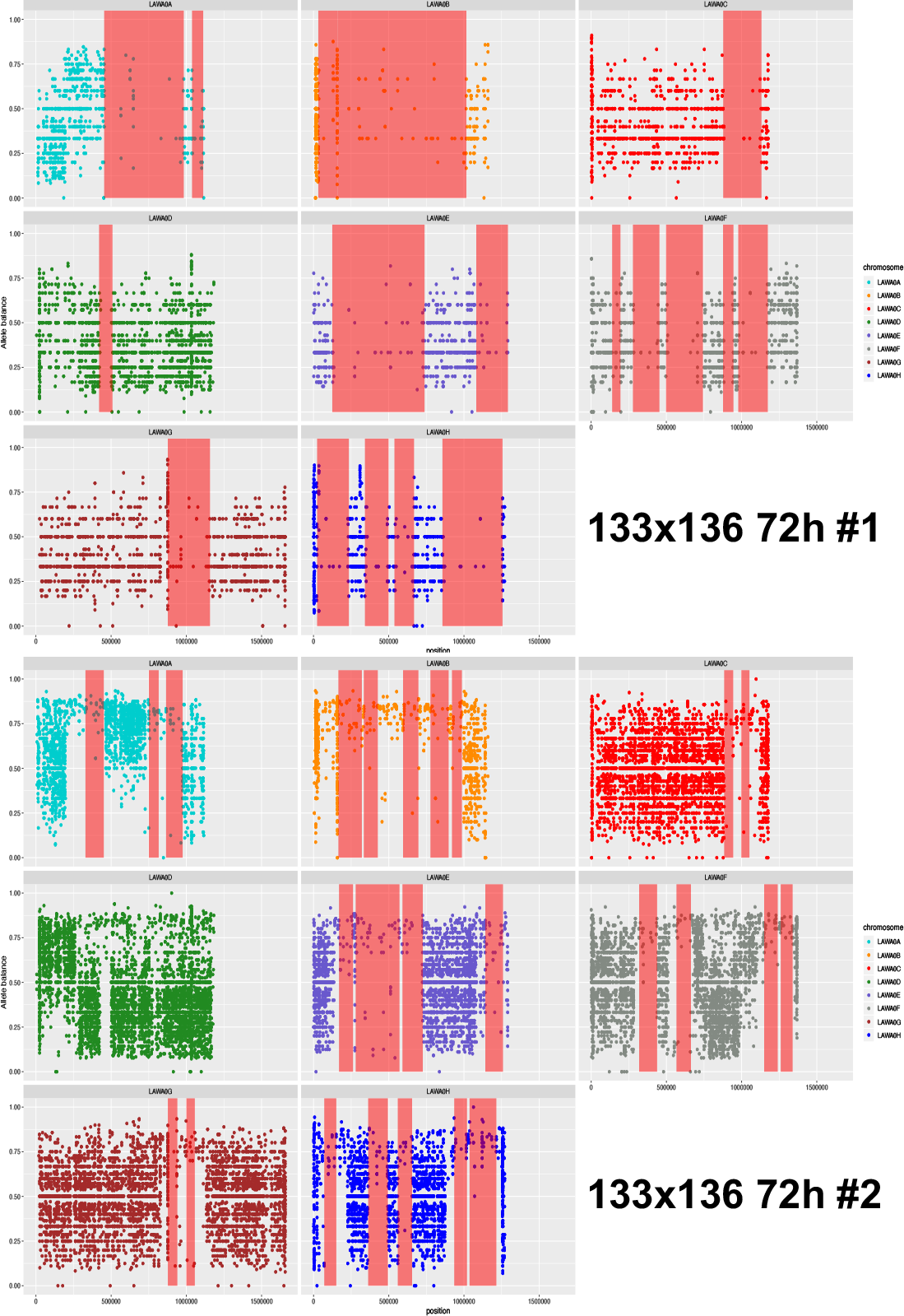
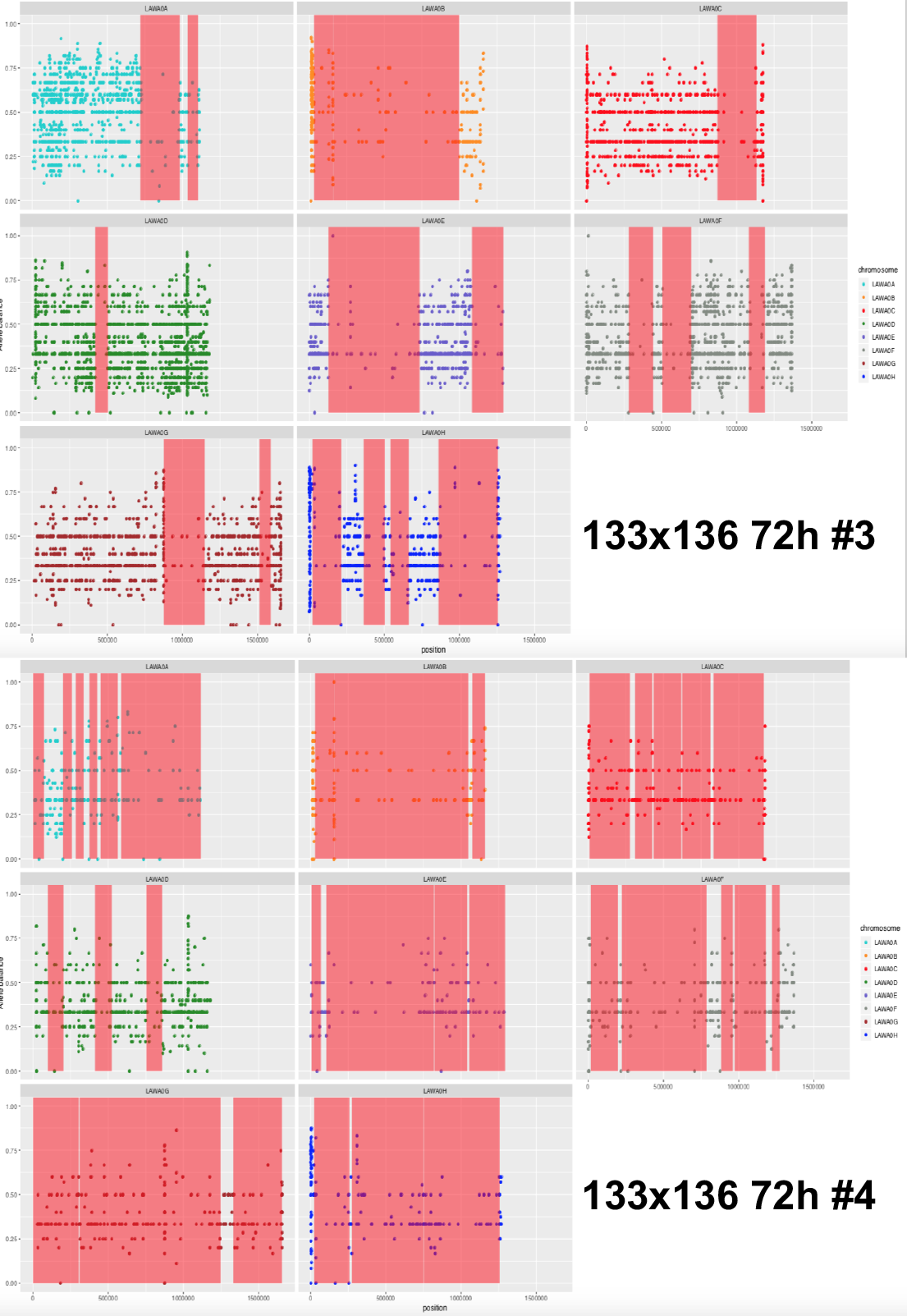
Distributions of LOH regions across the independent diploids generated from the LA128/136 (n=8) and LA133/136 (n=6) hybrids. The diploids were isolated at either 48 hours or 72 hours post mating from malt-agar plates. LOH regions (vertical red areas) were identified on the basis of the number of heterozygous sites per 50 kb windows as in Peter et al. 2018. Only heterozygous sites are represented on the different plots.

**Figure S4.**
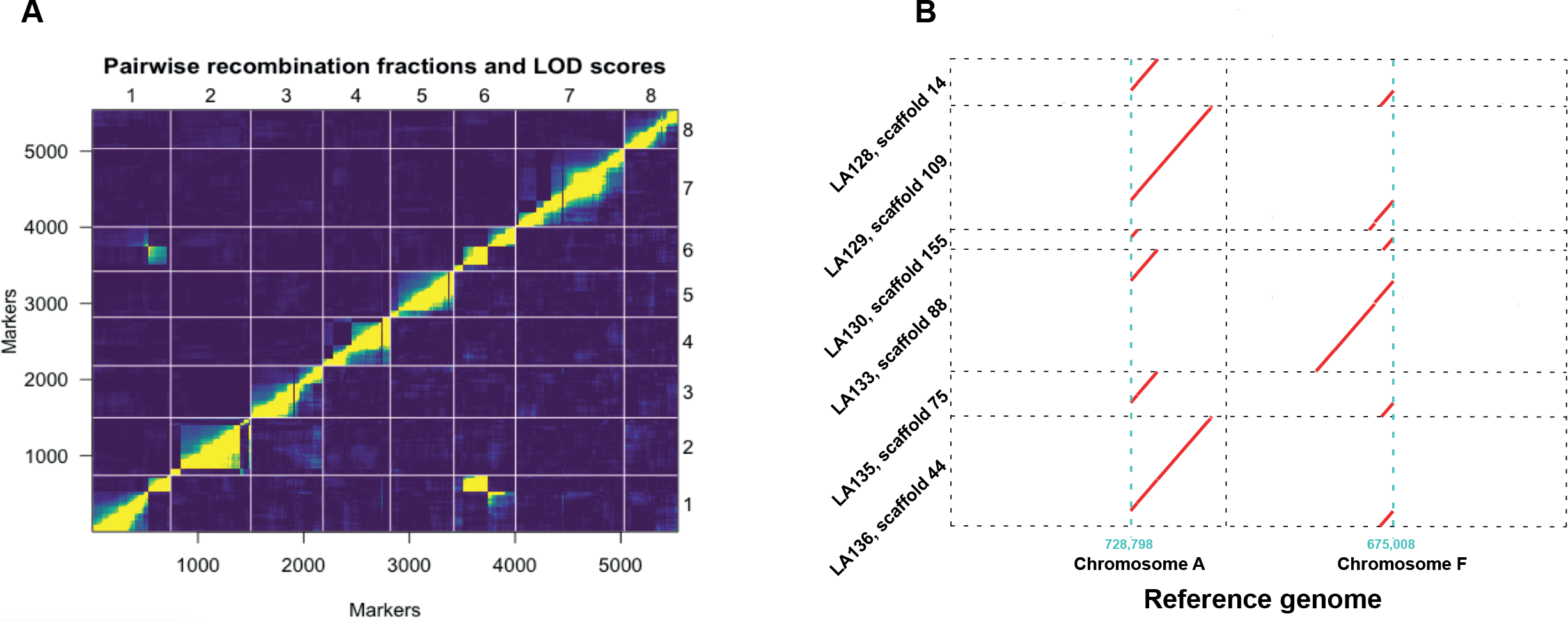
Reciprocal translocation between chromosomes A and F in LA128 and LA136 compared to the reference strain. **A**. Pairwise linkage disequilibrium for a representative subset of 5,542 markers. The markers in x-axis and y-axis are ordered according to their position along the reference genome and chromosomes are numbered from 1 to 8. Yellow represent high linkage disequilibrium and dark blue low linkage disequilibrium. Yellow spots between markers of the first and sixth chromosomes highlight the presence of a translocation in the progeny compared to the reference genome. **B**. Translocation between chromosomes A and F was detected in all Canadian isolates (728,798 on chromosome A and 675,008 on chromosome F).

**Figure S5.**
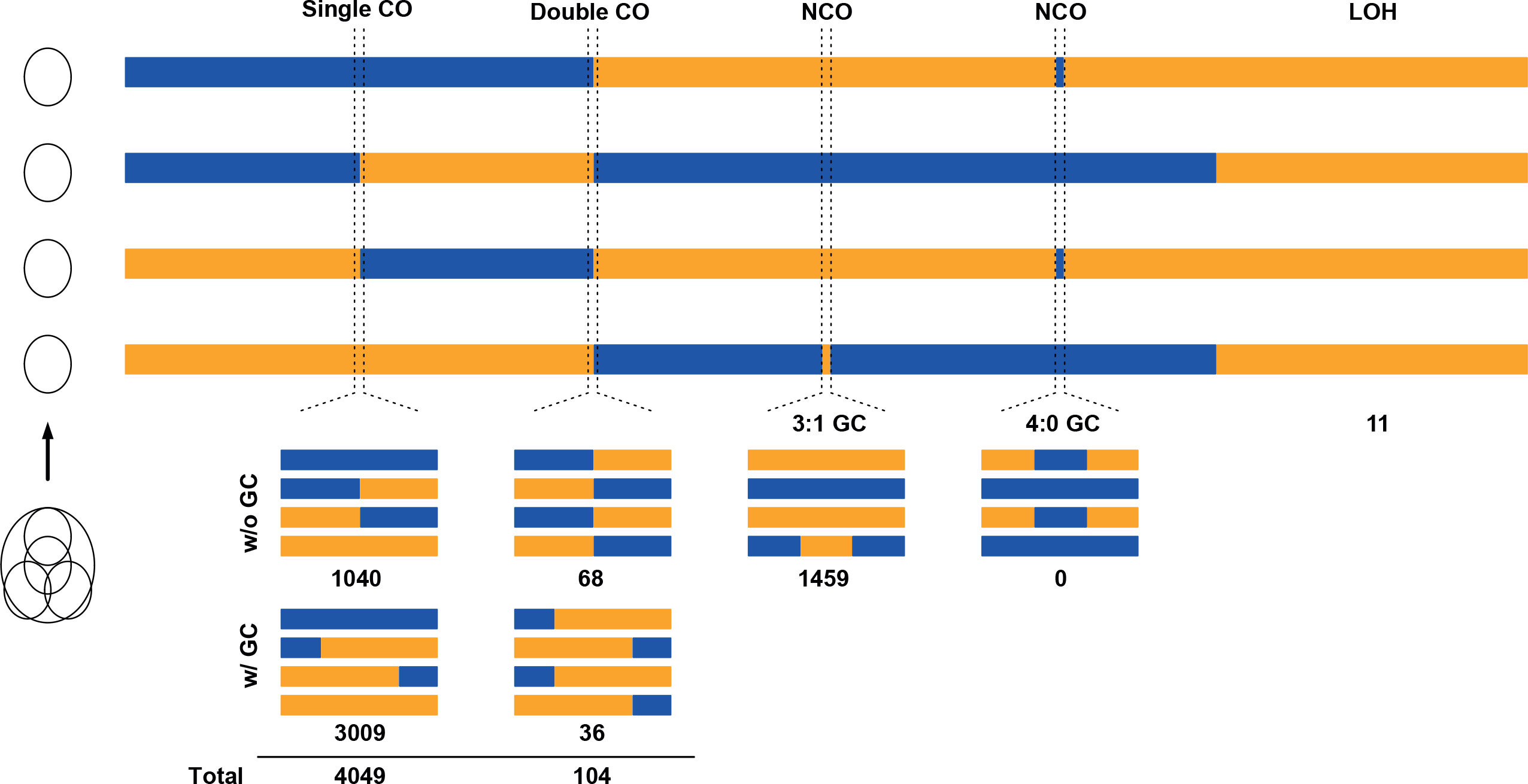
Allelic segregation patterns observed in the four spores of a tetrad. CO-Crossover, NCO-Non crossover, GC-Gene conversion, LOH-Loss of heterozygosity event during meiosis. Reported numbers are the counts of such events detected across the 192 tetrads.

